# Water chlorination increases the relative abundance of an antibiotic resistance marker in developing sourdough starters

**DOI:** 10.1101/2023.03.06.531128

**Authors:** Pearson Lau, Swapan Jain, Gabriel G. Perron

## Abstract

Multiple factors explain the proper development of sourdough starters. While the role of raw ingredients and geography, among other things, have been widely studied recently, the possible effect of water chlorination on the overall bacterial communities associated with sourdough remains to be explored. Here, using *16s rRNA* amplicon sequencing, we show that water chlorination at levels commonly found in drinking water systems has a limited impact on the overall bacterial communities developing in sourdough starters. However, using targeted sequencing, we found that the abundance of integron 1, a genetic mechanism responsible for the horizontal exchange of antibiotic resistance genes in spoilage and pathogenic bacteria, increased significantly with the level of water chlorination. While our results suggest that water chlorination might not impact sourdough starters at a deep phylogenetic level, they indicate that it can favor the growth of key spoilage bacteria.

## 1. Introduction

The use of sourdough starters is considered one of the great advancements in cooking (Chavan and Chavan, 2011). As the primary bread-leavening agent until the European industrial revolution (Pallant, 2021), sourdough starters increased the nutritional value of bread and made the latter a staple food in many parts of the world (Arora et al., 2021). More recently, sourdough regained popularity and has increasingly been celebrated for its desirable gastronomic properties (Arendt et al., 2007; Thiele et al., 2002)

Traditional sourdough starters are made from a mixture of flour and water fermented naturally by diverse populations of yeast and bacteria (Gobbetti, 1998). As the starter ferments, microbial populations that are initially diverse become quickly dominated by lactic acid bacteria and, to a lesser extent, by acetic acid bacteria (Arora et al., 2021; Ercolini et al., 2013). The dominant presence of lactic acid bacteria results in chemical, metabolic, and enzymatic activities that not only increase the nutritional value of sourdough bread but also inhibit the growth of other bacterial genera, contributing to the self-preserving properties of sourdough (Hansen and Schieberle, 2005; Liljeberg et al., 1995).

Multiple factors explain the proper development of microbial communities in sourdough starters (Landis et al., 2021). For example, sourdough starters are heavily influenced by temperature variations (Ercolini et al., 2013), and the ingredients used in starter generation (Reese et al., 2020; Ripari et al., 2016). However, the role of water, the other main ingredient in sourdough starter preparation, has only recently been recognized as a potential factor shaping microbial communities in sourdough starters (Minervini et al., 2019). More specifically, the presence of disinfectant residuals commonly used in drinking water is widely believed among professional bakers to negatively impact the proper development of microbial communities found in sourdough starter, potentially changing the flavor profiles of the sourdough (Leader and Blahnik, 1993; Vetri et al., 2020).

Chlorination is the most common disinfectant used in public water distribution systems (CDC, 2020). Most often, water chlorination is achieved by adding sodium hypochlorite, which leads to the presence of hypochlorite ion (OCl-) in the medium, inhibiting bacterial growth by disrupting metabolism and enzymatic inactivation (Estrela et al., 2002). The efficiency with which chlorination can stop the spread of most water-borne pathogens is considered one of the most outstanding achievements in public health of the past century (Li and Mitch, 2018). However, the inhibiting activity of chlorine present in water is not limited to pathogenic bacteria (Fish et al., 2020; Shi et al., 2013). Indeed, the potential impact of chlorine is predicted to extend to most microbial communities exposed to chlorinated water (Martino, 2019).

The presence of free chlorine in the water could alter the chemical properties of sourdoughs. For example, hypochlorite ion (OCl-) is an oxidizing agent that can break glycosidic bonds within bread starches, reducing the gluten network, and subsequently reducing the ability of flour components to gel together during the baking process (Egharevba, 2019). Water chlorination can also reduce flour’s lipid content due to the formation of chlorine derivatives (Bosmans et al., 2019). In addition to affecting the gustatory properties of the sourdough, such changes could affect developing microbial communities.

Chlorination was also demonstrated to promote the spread of antibiotic resistance (Jin et al., 2020; Zhang et al., 2022). While chlorination contributes to reducing the number of antibiotic-resistant bacteria in treated water at first (He et al., 2019; Ma et al., 2022), the continued presence of low concentrations of free chlorine in water can select for antibiotic-resistant bacteria downstream from treatment plants (Liu et al., 2018) and promote the exchange of antibiotic resistance genes among bacteria via horizontal gene transfer (Jin et al., 2020; Zhang et al., 2022). While the presence of antibiotic-resistance genes in sourdough and sourdough starters is unlikely to be a major health concern, even though DNA is not destroyed during cooking (James et al., 2021), antibiotic-resistance is often associated with spoilage bacteria (Samtiya et al., 2022). Therefore, selecting antibiotic-resistant bacteria in sourdough could affect the bread’s quality and preservation.

Here, using *16s rRNA* amplicon sequencing, we investigate the effect of chlorinated water on the development of bacterial communities in sourdough starters. In addition, we monitor the possible effect of water chlorination on the spread of integron 1, an important genetic element associated with the spread of antibiotic resistance in pathogenic bacteria (Gillings et al., 2015). While we show that water chlorination has a limited impact on the overall bacterial community structure developing in sourdough starters, we found that chlorinated water increased the abundance of integron 1, an indicator associated with clinically important antibiotic resistance genes.

## 2. Material and Methods

### 2.1 Establishment of sourdough starters

Sourdough starters are composed of two ingredients, flour and water, mixed together and regularly replenished to favor microbial growth. To control for possible variation in flour composition, we used a single bag of organic, stone-ground whole wheat flour (King Arthur Flour, Norwich, VT) for the entire experimental period of sourdough fermentation. According to the manufacturer’s website, the flour is made from dark northern hard red wheat, a varietal of the common wheat (*Triticum aestivum*) that contains a higher protein content (∼13.8%). We established the sourdough starters by combining 10 g of flour with 10 mL of control or treated water (see below) in sterilized polypropylene Nalgene bottles. Next, we mixed manually with an ethanol-sterilized glass rood, resulting in a dough yield (DY) of 200, or pastelike (Katina et al., 2014).

We then fed the sourdough starter every 24 hours (±2 hours), commonly called “backslapping,” by replacing 50% of the starter with a fresh mixture of flour and water. We ensured the complete homogenization of each sourdough starter by pouring the dough into a sterile bag and homogenizing it using the BagMixer 5000 (Interscience, Saint Nom la Brétèche, France) using default parameters for 60 seconds. We then remove 50% of the starter by weight and replace it with a fresh mixture of flour and water, as described above. Next, the dough was scraped down and remixed on the same settings. The freshly fed starter paste was then squeezed into a new sterilized polypropylene bottle. The bottles were placed in a dark cupboard for 24 hours at ambient room temperature maintained at 22-25°C. We repeated the feeding procedure six times for a total fermentation time of 7 days.

To identify the optimal growing conditions for investigating the possible effects of water chlorination, we established two independent trials. First, we chose to limit the exposure of the sourdough starters to bacteria in the air by having an “air-tight” container or screwing on the lid tightly. The only times the container lids were removed were to refresh the starter. Second, we conducted a second set of experiments with identical conditions, but this time allowing exposure to air and located in a working kitchen. For each experiment, we established three replicate starters for each control and chlorine treatment for a total of 18 experimental populations. All measurements and mixing were done under sterile, aseptic conditions throughout the study.

### 2.2 Water chlorination treatments

To test for the possible effects of water chlorination on sourdough starters, we established and maintained sourdough starters with three water chlorination treatments resulting in three concentrations of free chlorine in the water: 0 ppm (or control); 0.5 ppm (0.5 mg/L), and 4.0 ppm (4.0 mg/L). The treatments were chosen to reflect the minimum and maximum residual amount of chlorine in finished drinking water in the United States of America (CDC, 2014). We prepared chlorinated waters daily before sourdough feeding by diluting sodium hypochlorite, or NaOCl, into 100 mL of sterilized water.

We tested for the water chlorination level in each water preparation using the N, N-dimethylacetamide method (LaMotte, Baltimore, USA). Briefly, 5 mL of chlorinated water was mixed with N, N diethyl-p-phenylenediamine and was compared to the provided color chart, indicating the available chlorine concentration of the solution.

Because we were also concerned with the presence of other organic material in the water interfering with the disinfection efficacy of chlorine concentrations in our treatments, we tested the residual chlorine concentrations using the digital colorimeters method as described in the CDC protocol for measuring residual free chlorine (CDC, 2022). We confirmed that residual chlorine concentration was stable throughout our experiment.

### 2.3 Sample Processing

We used a *16S rRNA* amplicon sequencing approach to characterize the bacterial communities found in each sourdough starter, also known as food microbiomes. We first extracted the bacterial DNA from 1g of dough, which we diluted in 9 mL sterile, peptone physiological solution (0.1% peptone, 0.85% NaCl). We then extracted microbial DNA from the diluted sourdough using the procedure described in the MoBio PowerFood DNA Extraction kit (MoBio, Carlsbad, CA). Finally, we amplified the V4 region of the *16S rRNA* gene using the Golay-barcoded primers 515F and 806R (Caporaso et al., 2012).

Following gel purification, libraries were pooled at equimolar ratios and sequenced on the NANOseq paired-end Illumina platform adapted for 250-bp paired-end reads (Wright Labs, Huntingdon, PA). All unprocessed sequence reads are available at the Sequence Read Archive of the National Center for Biotechnology Information (NCBI accession number: PRJNA784321).

### 2.4 *Processing of* 16S rRNA *amplicon sequence data*

We characterized the microbiomes of each sourdough starter sample by identifying and tabulating the number of different sequence variants, also known as amplicon sequence variants (i.e., ASVs). Sequence variants can then be assigned to a taxonomic rank, usually at the genus level, providing additional information about the biology of each microbiome community. More specifically, we processed the *16S rRNA* reads using the *DADA2* pipeline version 1.20 (Callahan et al., 2016) available at (https://github.com/benjjneb/dada2) and implemented in *R* version 4.1.1 (http://www.r-project.org) (see Pindling et al. 2018 for full details) where each sequence read is assessed for contaminants, quality purposes, and possible sequencing error. Taxonomy was assigned using both the DADA2 native taxa identifier function as well as *IDTAXA* (Murali et al., 2018) available via the *DECIPHER* Bioconductor package (DOI: 10.18129/B9.2bioc.DECIPHER) trained on the SILVA ribosomal RNA gene database version 138.1 (Quast et al., 2013) as well as the RDP trainset 18 (McDonald et al., 2012; Werner et al., 2012). A complete list of all ASVs and their abundance in each sample can be found in Table S1, and a complete taxonomic assignment can be found in Table S2. Lastly, we build a maximum likelihood phylogenetic tree based on a multiple alignments of all the ASVs using the *phangorn* package version 2.1.3 (Schliep 2011); the latter is used to estimate the total phylogenetic, or evolutionary, distance present in each sample.

### 2.5 Microbial Community Analysis

Microbiomes’ diversity was analyzed using *phyloseq* version 1.30.0 (available at https://joey711.github.io/phyloseq/) implemented in *R* and visualized in *ggplot2* (Wickham, 2009). A mapping file linking sample names and the different treatments is provided in Table S3. Briefly, for each sample, we estimated the total number of ASVs, Chao1, which is the predicted number of ASVs in the whole sample, as well as diversity as Simpson’s Index, i.e., *D*, from the ASV table subsampled to the lowest sampling depth adjusted for each comparison. While richness considers the total number of ASVs, diversity includes evenness measures among the different ASVs present in a sample. We tested whether richness and diversity indices differed among treatments using linear modeling and comparing the different statistical models with Akaike’s Information Criterion (AIC) as implemented in *R’s* stats package.

To test whether there were statistical differences in population structure between treatments (e.g., controls microbiomes vs. microbiomes exposed to chlorinated water as well microbiomes exposed to air vs. microbiomes not exposed to air), we performed Principal Coordinate Analyses (PCoA) on phylogenetic distances calculated as weighted UniFrac distance scores (Lozupone and Knight 2005). We used a permutational analysis of variance (PERMANOVA) implemented via the *adonis* function (Oksanen et al., 2015) of *vegan* version 2.5.6 to test for significance. Finally, we confirmed that each test respected the homogeneity of variances assumption using the betadisper method of the *vegan* package (Oksanen et al., 2015).

### 2.6 Quantify the Presence of an Antibiotic Resistance and Spoilage Bacteria Marker

Finally, we investigated whether chlorinated water increased the relative abundance of *intI1*, a gene encoding integron class 1 integrase (Gillings et al., 2015, p. 1). The latter is almost entirely associated with spoilage or potentially harmful bacteria and facilitates the spread of antibiotic-resistance genes among bacteria (Gaze et al., 2011; Zhang et al., 2018). As described elsewhere (Gaze et al., 2011), we quantified *16S rRNA* and *intI1* gene copy numbers from triplicates reactions using the Bio-Rad CFX96 Real-Time PCR Detection System (Bio-Rad Laboratories, Hercules, CA, USA). An internal standard curve was included with each qPCR run. We then calculated the relative abundance of the *int1* by normalizing its copy number to that of the *16S rRNA* copy number in each sample. The latter was adjusted by dividing it by 4.2, the average number of *16S rRNA* copies in each bacteria cell (Větrovský and Baldrian, 2013). Finally, we used linear modeling to test for the effect of water chlorination on *intI1* relative abundance and compared the different statistical models with Akaike’s Information Criterion (AIC) as implemented in *R’s* stats package. While we used the square root-transformed data for statistical analysis, we plotted the raw data.

## 3. Results

### 3.1 Overall Microbial Population Structure

We characterized the microbiomes of 36 sourdough starters by sequencing the V4 region of the 16S *rRNA* gene. Excluding eight samples that failed to be sequenced, we obtained a total of 1,160,000 pairs of forward and reverse reads with an average read length of 250 base pairs, totaling ∼583 G bases and an average sequencing depth per sample of 41,642.9 paired-reads. After trimming forward reads at 240 bp, reverse reads at 225 bp, and removing predicted *phiX* and chimeric reads, we were left with 980,261 (84.1% of the initial) paired-reads.

We identified a total of 150 unique ASVs with a median of 9 ASVs found in each starter (Table S1). The number of ASVs found in each starter varied greatly from 4 ASVs found in a starter fermented for 1 day to up to 65 ASVs in a starter fermented for 7 days. In total, ASVs were matched to 82 different bacteria genera (Table S2), with *Latilactobacillus* being the most common among our samples, 49.4(39.5)%, followed by *Pantoea* sp., 27.2(39.0)%, *Weissella* sp., 9.1(23.2)%, and *Pseudomonas* sp., 8.7(26.7)%. As taxonomy changed slightly depending on the database and algorithm used (Figure S1), we will present our results using the RDP database with the *TaxaID* taxonomy identifier thereafter; the latter offered the smallest number of unidentified ASVs.

### 3.2 Identifying the source of microbial fermentation in new sourdough starters

We first wanted to identify the ideal condition to establish sourdough starters under our laboratory conditions. To do so, we established two sourdough starters trials: one under strict laboratory sterile conditions and limited exposure to air and the second in kitchen environments with exposure to air. When comparing the microbial communities found in the sourdoughs after 7 days of fermentation and using the same feeding procedure, we found a significant divergence in the population structure (ADONIS: *F*_(1,16)_ = 27.7; *R*^2^ = 0.65; *P* = 0.001; Figure 1). The main difference we observed between the two trials was that the most common ASV found in the presence of air was identified as *Latilactobacillus* sp.. In contrast, the most common ASV found under sterile conditions was identified as *Panteoa* sp., a genus not commonly associated with sourdough starters. Furthermore, while we did not find significant differences in the number of observed ASVs (*F*_(1,15)_ = 5.08; adj-*P* = 0.16; Figure S2A) or evenness measured as the Simpson’s Index (*F*_(1,25)_ = 4.16; adj-*P* = 0.18; Figure S2B), we found that the starters grown in sterile conditions were more prone to invasion, as defined by a microbial community where a lactic acid bacteria did not represent at least 80% of the total identified ASVs (Fishers Exact test: *P* = 0.05). For the above reasons, we decided to investigate the possible effect of water chlorination on sourdoughs exposed to air only.

**Figure 1.**
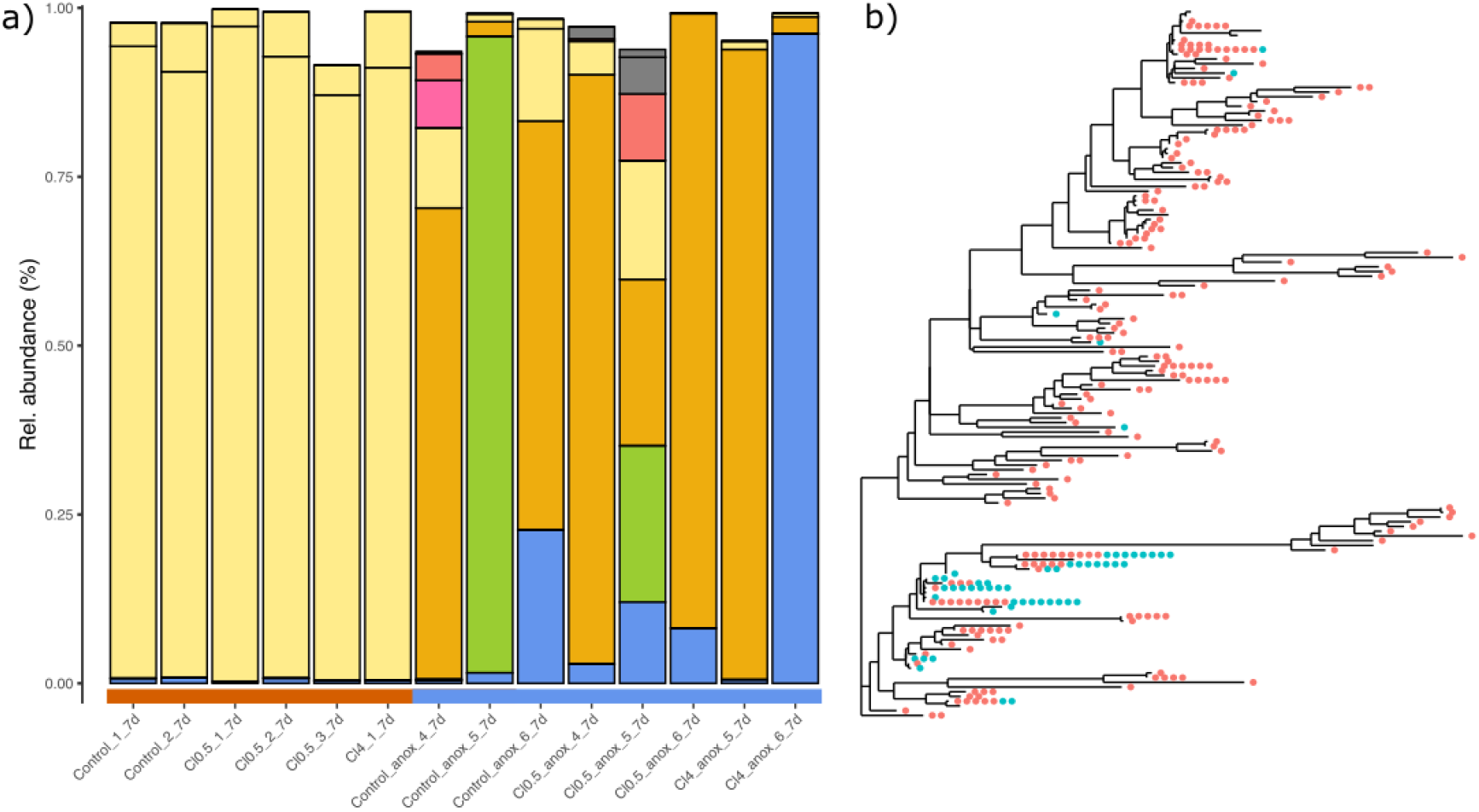
Microbial diversity found in sourdough starters in unfiltered air (red) and filtered air (blue. (a) Relative abundance of the eight most common bacteria genera: *Enterococcus* (green), *Klebsiella* (pink), *Pantoea* (dark gold), *Latilactobacillus* (heirloom corn), *Pseudomonas* (olive), *Erwinia* (red), *Leuconostoc* (red), *Weissella* (blue). (b) Phylogeny of all ASVs represented as a maximum likelihood tree shows that bacteria types found in sourdough fermented in the presence of unfiltered and filtered air were more clustered than expected by chance alone. Sourdough starters fermented in fileted air are shown in blue, while starters fermented in unfiltered air are presented in red.

### 3.3 The effect of water chlorination on sourdough starters

To test for the possible effect of water chlorination on the microbial communities developing in sourdough starters, we exposed starters to three different chlorination treatments, including a control group not exposed to hypochlorite ion. We found no evidence that water chlorination affected the overall structure of microbial communities in sourdough starters. Overall, the same few ASV dominated the populations by day 7 in all treatments (Figure 2A). Using principal component analysis to detect possible changes in community structure measured as phylogenetic distance via Unifrac scores, we found that microbial populations changed significantly over time (ADONIS: *F*_(1,15)_ = 8.01; *R*^2^ = 0.35; adj-*P* = 0.004; Figure 2A) similarly in all water chlorination treatments (ADONIS: *F*_(2,14)_ = 0.12; *R*^2^ = 0.02; adj-*P* > 0.99; Figure 2B). In other words, even a chlorine concentration at a level observed in some of the most chlorinated public water systems, i.e., 4 ppm, did not modify the relative abundance of most ASVs and taxa observed in the different samples.

**Figure 2.**
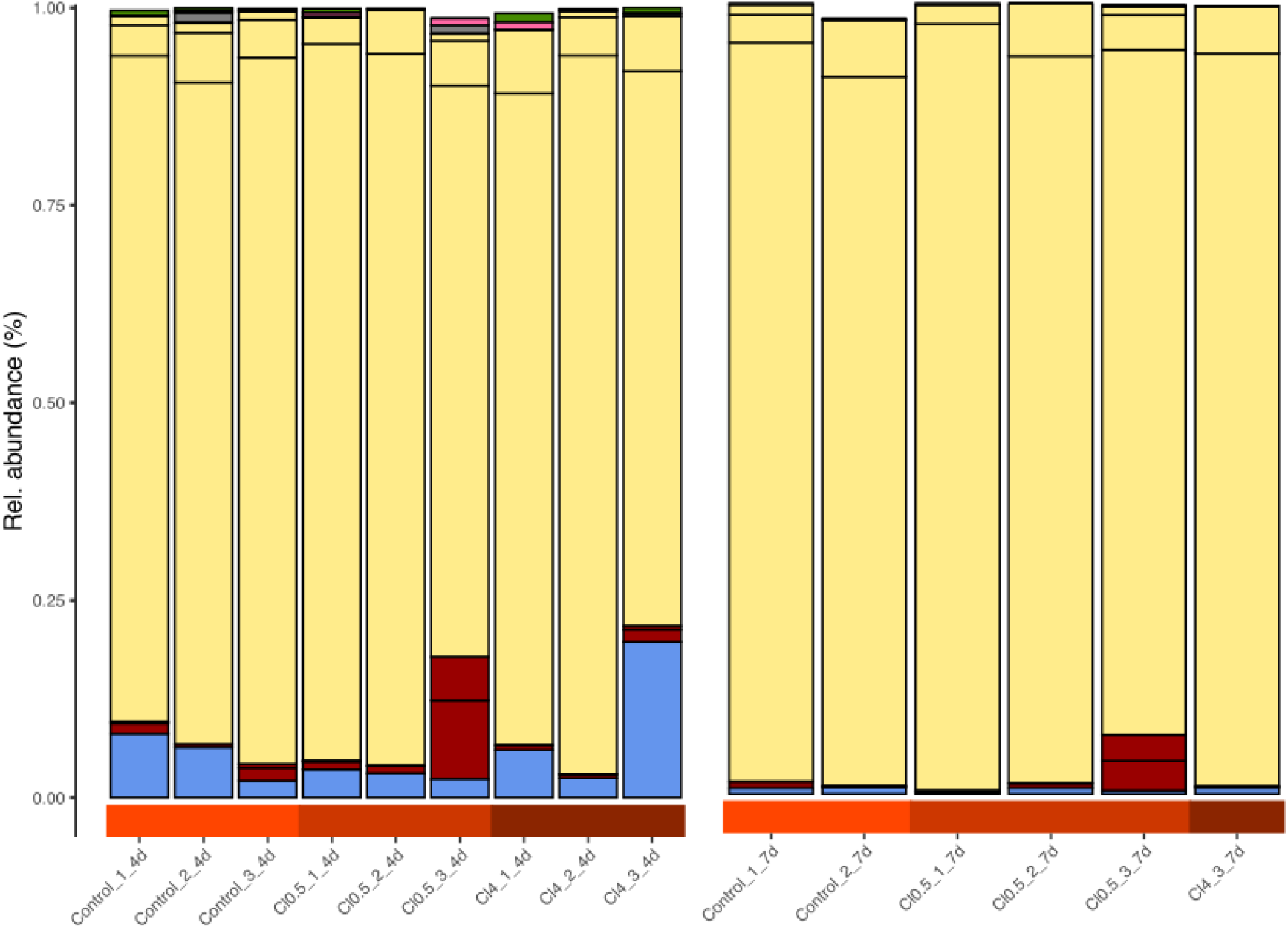
Effect of water chlorination on sourdough starter microbial communities. Relative abundance of the eight most common bacteria genera: *Enterococcus* (green), *Klebsiella* (pink), *Pantoea* (dark gold), *Latilactobacillus* (heirloom corn), *Pseudomonas* (olive), *Erwinia* (red), *Leuconostoc* (red), *Weissella* (blue). Taxonomy was identified at the genus level using the RDP training dataset v18 and the TAXAID function of the *DECIPHER* package as implemented in *R*. Concentration of chlorine in water tested was 0 μg/L shown in light red, 1 μg/L in red, and 4 μg/L in dark red.

Similarly, while the number of ASVs observed in the starters decreased over time (*F*_(1,5)_ = 15.32; *P* = 0.01; Figure 3A), we found that the number of ASVs did not differ between the water chlorination treatments (*F*_(2,5)_ = 0.10; *P* = 0.904; Figure 3A). We found similar results when considering the predicted total number of observed ASVs using the Chao1 index (*F*_(2,5_ = 0.10; *P* = 0.90; Figure S3A). Finally, we found that chlorine levels did not affect diversity as measured by Simpson’s index (*D*), a measure that is sensitive to how evenly the ASVs are distributed in the samples (*F*_(2,5_ = 0.02; *P* = 0.98; Figure S3B). Similarly to observed ASVs, however, diversity decreased over time (*F*_(1,5_ = 11.92; *P* = 0.02). Interestingly, when we look at the dispersion in the number of ASVs around the mean for each treatment, as measured by variance, we note that variance seems to increase in the presence of chlorine. Unfortunately, we do not have enough data points to test for this pattern. Still, this result suggests that water chlorination could result in finer changes while not changing the core microbial communities in the starters.

**Figure 3.**
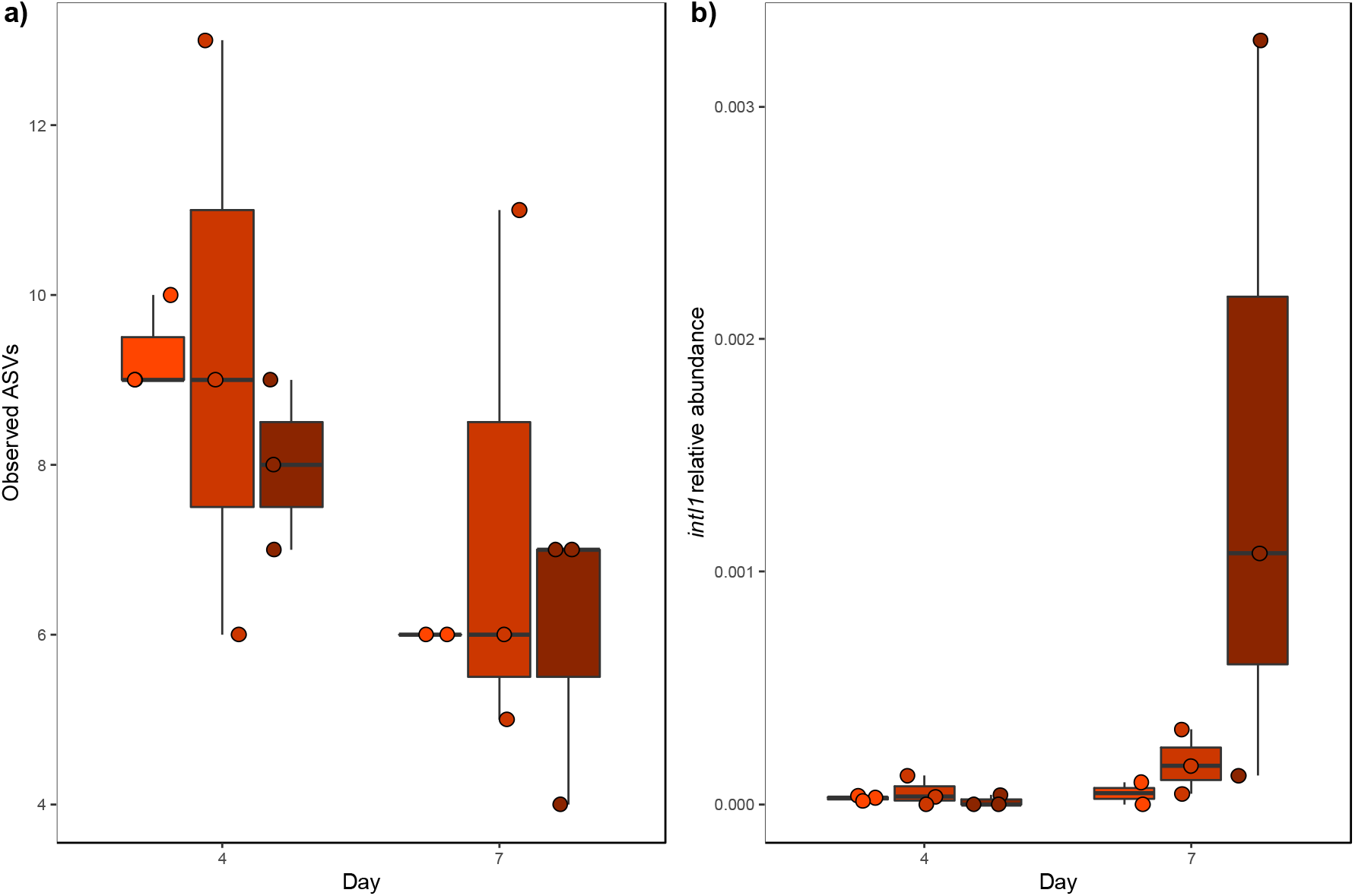
Effect of water chlorination on a) bacterial diversity and b) the relative abundance of intI1 in sourdough starters. A) Chlorination did not affect bacterial diversity measured as the number of observed ASVs (*F*_(2,5)_ = 0.10; *P* = 0.904), but affected B) *intI1*’s relative abundance, calculated as the proportion of *intI1* copy numbers per *16S rRNA* copies (treatment:time: *F*_(2,18)_ = 4.17; *P* = 0.03). Concentrations of chlorine in the water tested were 0 mg/L, shown in light red, 1 mg/L in red, and 4 mg/L in dark red.

### 3.4 Investigating the presence of integron 1 during fermentation

While we did not observe major changes in community structure or diversity, our previous results suggest that changes in community dynamics could have happened at a finer scale. For example, it is possible that chlorine could exert selective pressures on a resistant strain within a genus or even at the gene level via horizontal gene transfer. For this reason, we investigated the relative abundance of *intI1*, the gene encoding for integron class 1. The latter is a genetic mechanism enabling the quick transfer of genes and is almost always associated with bacteria with a spoiling or pathogenic potential and is usually associated with antibiotic resistance (Ghaly et al., 2021; Gillings et al., 2015; Zhang et al., 2018).

Using quantitative PCR, we found that chlorinated water affected the relative abundance of *intI* in sourdoughs over time (treatment:time: *F*_(2,18)_ = 4.17; *P* = 0.03; Figure 3B). More specifically, we found that the highest chlorine concentration, i.e., 4 ppm, significantly increased the relative abundance of *intI1* by day seven (*t* = 2.59; *P* = 0.02; Figure 3A). Interestingly, we found no difference in *intI1* relative abundance among the different chlorine concentrations at day four. In other words, a larger proportion of the bacteria detected via qPCR harbored the *intI1* gene by the end of the experiment only at the highest chlorine concentration. In contrast, the relative abundance of the gene stayed more or less constant across all other treatments over time.

## 4. Discussion

Multiple factors, such as flour quality or fermentation time, explain the proper development of sourdough starters (Ganzle and Ripari, 2016; de Vuyst et al., 2014; Landis et al., 202(Minervini et al. 2014). Understanding the factors that influence and shape the development of sourdough starters is not only important for the reliable production of quality sourdough but also can help us shed new light on the cultural importance of bread making. Here, we show how air and water chlorination can influence the development of microbial communities found in sourdough starters. We also found that sourdough starters affected by a chemical stressor like free chlorine present in water can be less resilient to the presence of possible food-spoiling bacteria or pathogenic bacteria.

More specifically, we found that sourdough starters exposed to unfiltered air developed healthy microbial communities with dominant bacteria taxa most often associated with traditional sourdough fermentation. On the other hand, when sourdough starters were exposed to filtered air in a controlled laboratory setting, the starters were dominated by *Panteoa* sp., an Enterobacteriaceae which was initially isolated as a plant pathogen and that can also be isolated from human and animal gut as well as spoiled soil and water (Walterson and Stavrinides, 2015). Some species of *Panteoa* are known contaminants of sourdough and can negatively affect fermentation (Celano et al., 2016; Ercolini et al., 2013). The fact that *Pantoea* dominated all of our starter replicates grown independently suggests that the possible contaminant was likely present in the flour we used to establish the starters and that exposure to unfiltered air is crucial for the proper development of healthy sourdough starters.

Interestingly, the amount of chlorine present did not affect the overall microbial community structure in healthy sourdough starters. Regardless of the chlorine concentration used, the same dominant bacteria taxon, i.e., *Latilactobacillus* sp., was detected in all starters by the end of the experiment. The latter is a lactic acid bacterium commonly identified in sourdough starters (Baek et al., 2021; Xu et al., 2020) and other fermented products (Liang et al., 2020; Liu et al., 2020). In fact, *Latilactobacillus* sp. accounted for more than 80% of the total read count in all our sourdough starters, confirming that chlorine did not affect our ability to produce healthy sourdough starters as previously predicted.

Our results show that chlorine could affect microbial communities at a finer scale. We found that the relative abundance of the gene *intI1* increases significantly with chlorine concentration in water. In other words, the relative abundance of *intI1* was significantly higher in starters exposed to the highest free chlorine level when compared to the other treatments. While the copy number of *intI1* was relatively low compared to the total number of *16S rRNA* gene copies identified in our study, the presence of a marker associated with antibiotic-resistance genes and spoilage bacteria should be taken seriously. Whether our observation of this gene marker translates into the actual presence of antibiotic-resistant bacteria in sourdough remains to be tested. Yet, it is known that some bacterial strains associated with sourdough fermentation show intrinsic resistance to antibiotics (Manini et al., 2016), and that farming systems where raw ingredients were grown can also contribute to the presence of antimicrobial resistance composition (Rizzello et al., 2015). Finally, even if the genomic content of starter cultures does contain a high abundance of antibiotic-resistance genes, how it affects subsequent functionality in fermented food has yet to be determined (Leech et al., 2020).

In conclusion, our study provides an important proof-of-principle of the possible effect of water chlorination on sourdough starters and contributes to the growing body of literature investigating how environmental variables shape fermented foods. Our findings also suggest that whole genome sequencing and proteomics might be required to fully understand the finer changes in microbial communities impacted by water chlorination and other environmental factors, possibly impacting the desired gastronomic properties of sourdough bread.

## Acknowledgments

This work did not receive any specific grant from funding agencies in the public, commercial, or not-for-profit sectors. The authors would like to thank Maureen O’Callaghan-Scholl and Chef Rei Peraza for their technical assistance in the laboratory and the kitchen.

## Supplementary Materials

**Figure S1.**
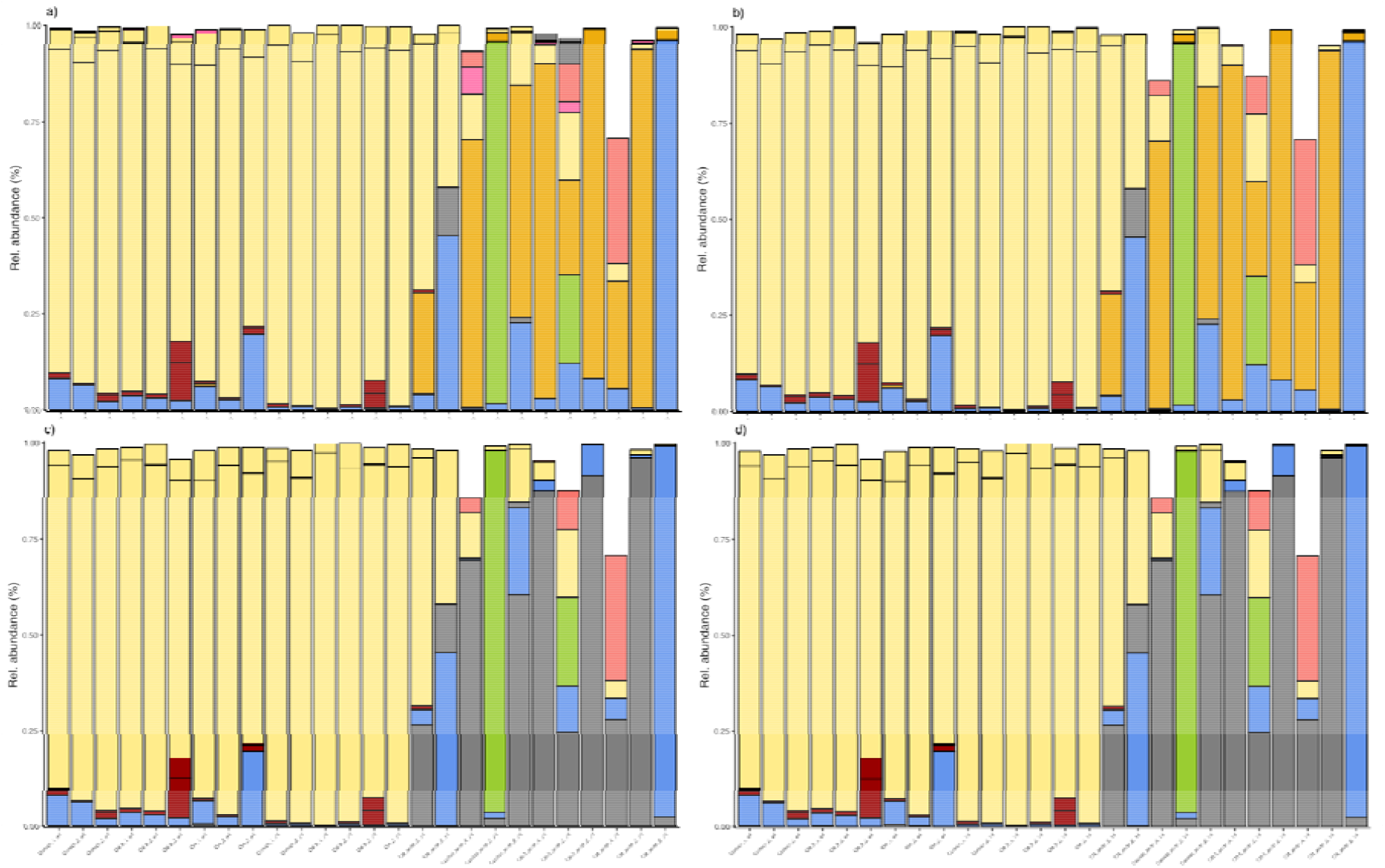
The effect of database and algorithm choice on taxonomic identification of the most common bacteria variants. Relative abundance of the top ten genera identified using the SIVLA training dataset v138.1 with a) the IDTAXA function from the *DECIPHER* package, and b) the *DADA2* native “assignTaxonomy” function as well as the RDP training dataset v18 with c) the IDTAXA function from *DECIPHER*; and d) the *DADA2* “assignTaxonomy” function. Genera are identified as follow: *Enterococcus* (chartreuse), *Klebsiella* (pink), *Lactococcus* (dark goldenrod), *Latilactobacillus* (light goldenrod), *Leuconostoc* (plum), *Weissella* (blue).

**Figure S2.**
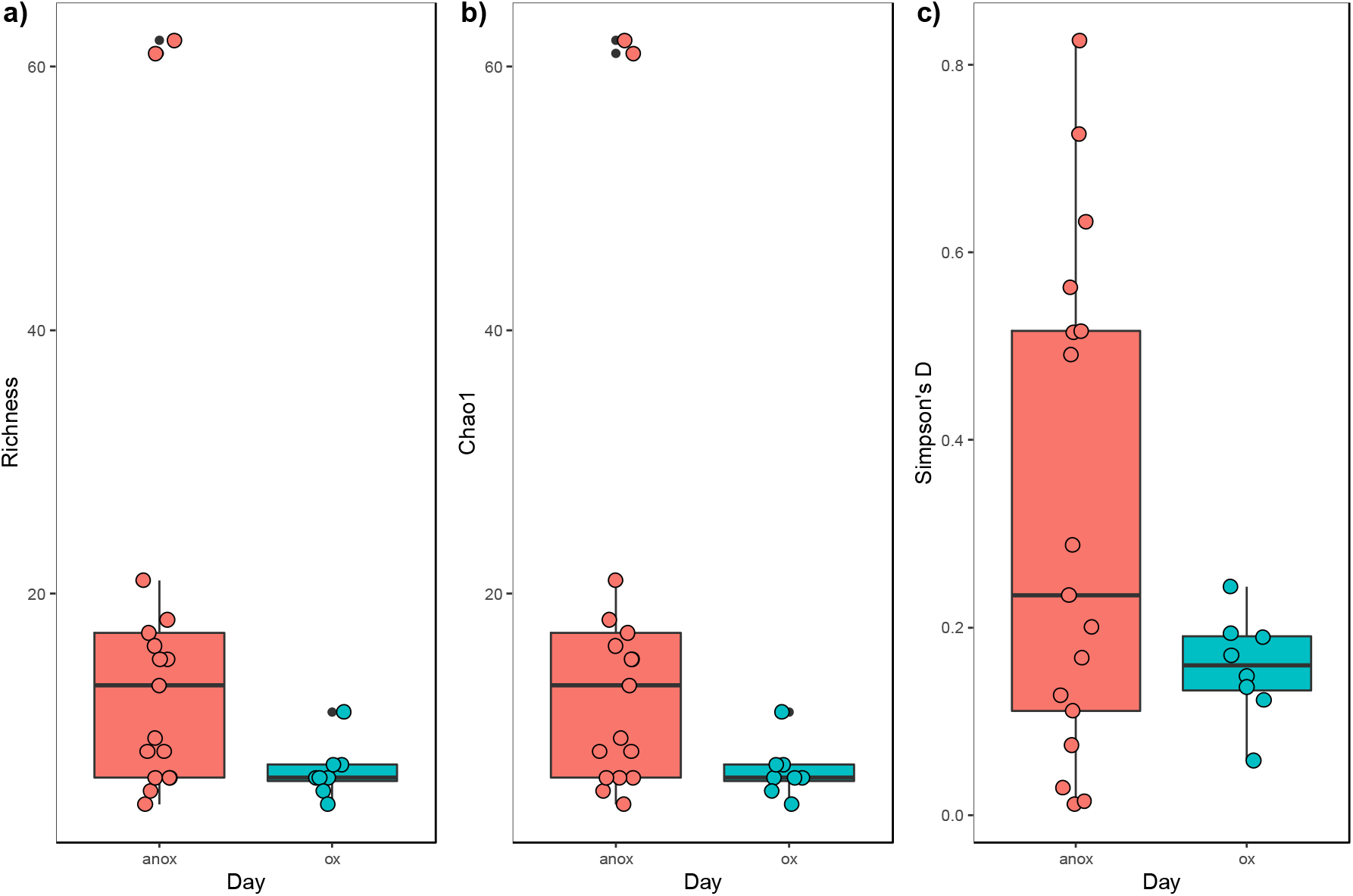
Effect of exposure to air on a) bacterial diversity and b) the relative abundance of intI1 in sourdough starters. A) Bacterial diversity is measured as the number of observed amplicon sequence variants (or ASV) and B) Chao1, as well as C) Simpson’s D. Starters grown in the presence of air are shown in blue and starters grown in anaerobic conditions are shown in red.

**Figure S3.**
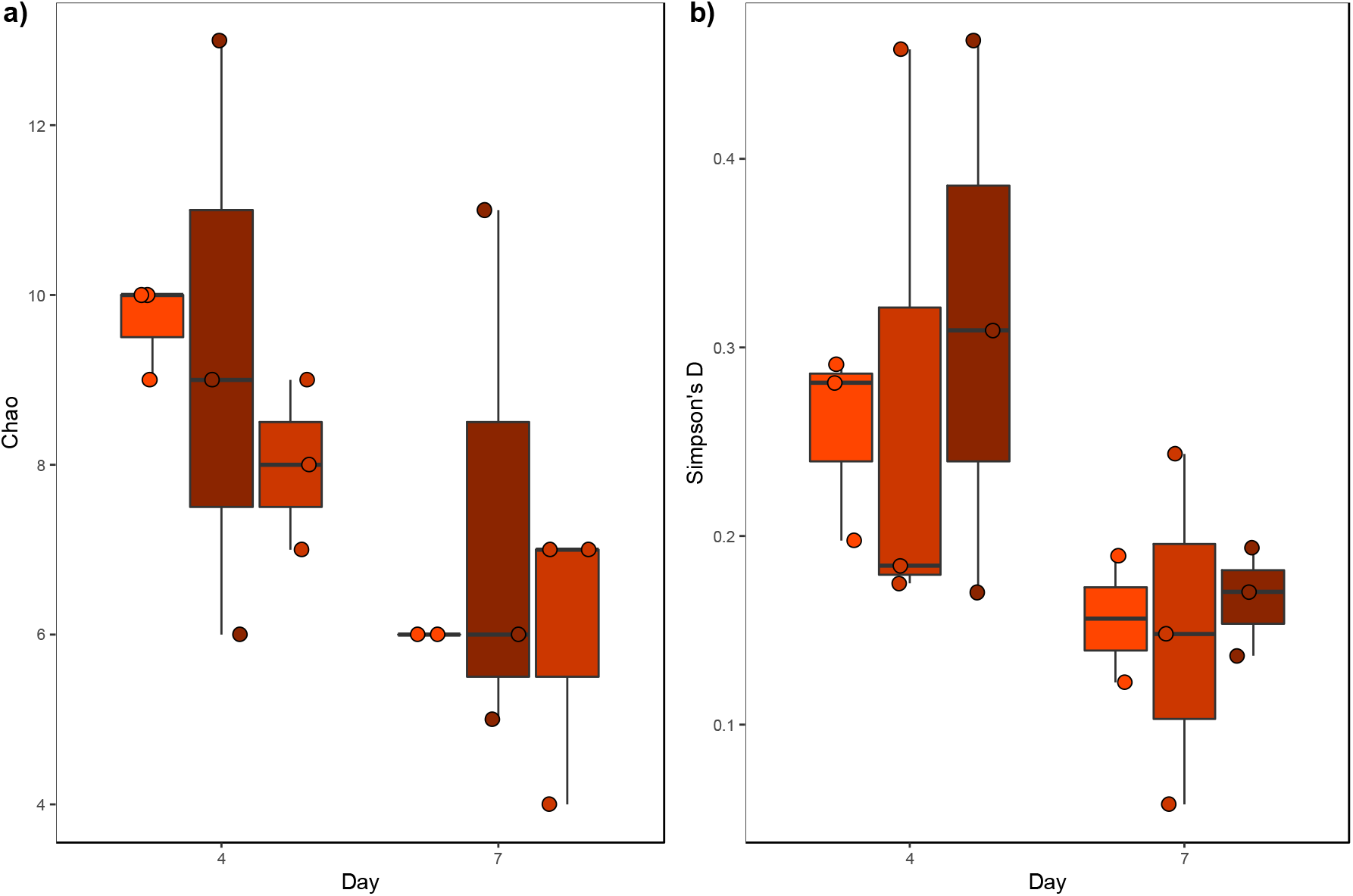
Effect of water chlorination on bacterial communities measured as a) predicted observed ASVs and b) diversity. A) Bacterial richness is measured as Chao, or the number of predicted ASVs. B) Diversity is measured as Simpson’s D. Concentrations of chlorine in the water tested were 0 mg/L, shown in light red, 1 mg/L in red, and 4 mg/L in dark red.

